# Enhanced γ-globin reactivation and sickle cell correction through a repressor-to-activator motif switch in the *HBG1/2* promoters

**DOI:** 10.64898/2026.04.07.716887

**Authors:** Anne Chalumeau, Panagiotis Antoniou, Maria Bou Dames, Mike Firth, Martin Peterka, Marcello Maresca, Mégane Brusson, Annarita Miccio

## Abstract

Sickle cell disease (SCD) is caused by the production of an abnormal adult hemoglobin that generates sickle-shaped red blood cells (RBCs). Transplantation of autologous genetically corrected hematopoietic stem/progenitor cells (HSPCs) represents a promising therapy. Persistent fetal hemoglobin expression improves SCD.

Here, we engineered the fetal *HBG1/2* promoters by replacing the BCL11A repressor binding site (BS) with a TAL1:GATA1 motif recognized by transcriptional activators.

We exploited the prime editing nuclease (PEn) that efficiently installed the TAL1:GATA1 motif in K562 cells, outperforming the original PE. Non-homologous end joining (NHEJ) and/or alternative-end joining (alt-EJ) pathway inhibition enhanced precise editing. However, this strategy was poorly efficient in patients’ HSPCs.

Alternatively, we used CRISPR/Cas9 nuclease to either disrupt the BCL11A BS via NHEJ and/or alt-EJ or to replace it with the TAL1:GATA1 motif via homology-directed repair (HDR) using a donor ssODN template. NHEJ and alt-EJ inhibition improved product purity, reducing InDels and achieving superior precise editing efficiency compared to PEn in K562 and HSPCs. HDR-edited HSPCs preserved clonogenic capacity and differentiated into RBCs showing elevated *HBG* expression and correction of the sickling phenotype.

These results demonstrate that replacing the BCL11A BS with a TAL1:GATA1 motif is a potent strategy for reactivating *HBG1/2* to treat SCD.

## INTRODUCTION

Beta-hemoglobinopathies are genetic disorders caused by mutations that either reduce adult β-globin chain production (β-thalassemia) or generate an abnormal β-globin chain (sickle cell disease, SCD). Beta-thalassemia and SCD are the most common inherited genetic disorders affecting millions of people worldwide. In β-thalassemia, reduced β-globin synthesis leads to inadequate hemoglobinization of red blood cells (RBCs) and anemia. In SCD, a single point mutation in the β-globin (*HBB*) gene generates the sickle β^S^-globin chain and hemoglobin S (HbS) that polymerizes under hypoxic conditions. HbS polymerization triggers RBC sickling, ultimately resulting in vaso-occlusive crises, hemolytic anemia and organ damage. Current symptomatic treatments, such as RBC transfusions and supportive care, are associated with high costs, and continue to result in a poor quality of life. A curative option is allogeneic transplantation of hematopoietic stem and progenitors cells (HSPCs), but this is limited by immunological risks and availability of compatible donors^1^.

The clinical severity of β-hemoglobinopathies is alleviated by the co-inheritance of genetic mutations termed hereditary persistence of fetal hemoglobin (HPFH), which promote continued expression of the fetal γ-globin and production of fetal hemoglobin (HbF) in adult life^2^. Gamma-globin compensates for β-chain deficiency in β-thalassemia and exerts an anti-sickling effect in SCD^3^. Naturally occurring HPFH mutations identified in the promoters of the two γ-globin genes (*HBG1/2)* are known to either generate *de novo* DNA motifs recognized by potent transcriptional activators (e.g., KLF1^4^, TAL1^5^ and GATA1^6^) or disrupt/delete transcriptional repressor (e.g., LRF^7^ and BCL11A^7^) binding sites (BSs). Interestingly, the co-occurrence of multiple HPFH mutations is associated with higher HbF levels compared to individual mutations^8^.

Several genome editing strategies have been developed to genetically modify autologous HSPCs and treat patients with β-hemoglobinopathies. Most of these approaches are based on the use of a single-guide RNA (sgRNA) that drives the *SpCas9* (Cas9) nuclease to the target sequence where it generates a double-strand break (DSB). The DSB is then mainly repaired via the non-homologous end joining (NHEJ)^9^ or alternative-end joining (alt-EJ, also called micro-homology-mediated end joining, MMEJ) pathways^10,11^ leading to the generation of small insertions and deletions (InDels). This can be exploited to knock-out genes or inactivate regulatory elements. CRISPR/Cas9 has been used to inactivate the erythroid enhancer of *BCL11A,* a gene encoding a major HbF repressor^12,13^, showing promising results in pre-clinical and clinical trials. Exagamglogene autotemcel became the first FDA-approved CRISPR-Cas9 strategy to treat patients with β-hemoglobinopathies^14,15^. However, *BCL11A* knock-down can affect erythropoiesis, prompting the search for more suitable therapeutic targets^16,17^. CRISPR/Cas9 has also been used to reactivate HbF by disrupting the LRF repressor BS^18,19^ or the BCL11A BS^18,20–22^ in the −200 region and the −115 region of the *HBG1/2* promoters, respectively, via generation of InDels (predominantly deletions) through NHEJ or alt-EJ. In particular, approximately 30-50% of these deletions are associated with microhomology (MH) motifs at the target site, suggesting DNA repair via alt-EJ^23^. Precise genome editing can be achieved upon Cas9-mediated DSBs via homology-directed repair (HDR) in the presence of a DNA donor template. Cas9-HDR approaches are less efficient than NHEJ-based strategies in quiescent HSPCs; however, several optimizations have been carried out to increase Cas9-HDR efficiency, such as the use of NHEJ pathway inhibitors^24,25^.

Prime editing is a recently developed technology enabling all twelve possible base conversions, as well as targeted insertions, deletions, or combined edits^26^ in a specific region without generating DSBs. Prime editing utilizes a prime editor (PE) - a Cas9 nickase (Cas9n) fused to an engineered reverse transcriptase (RT)- together with a prime editing guide RNA (pegRNA) containing both a RT-template (RTT) with the desired edits and a primer binding site, which allows direct incorporation of edits into genomic DNA (PE2). To improve processivity and editing activity, the enzyme has been codon optimized and engineered by adding nuclear localization signal (NLS) sequences, linkers, and the R221K and N394K mutations in the Cas9n to improve its activity, generating the PEmax^27,28^. Despite all these improvements, some target regions remained hard-to-edit with the nickase-based PE, especially when DNA insertions are desired. We previously showed that this limitation can be overcome by replacing the nickase with a Cas9 nuclease, generating PEnmax, which supports precise insertion of DNA sequences and improves editing efficiency for pegRNAs that were inefficient with the nickase-based PE^29^. Of note, in the PEn-based strategy, a 30-nt long homology tail is added to the 5’ extremity of the RTT of the pegRNA to favor edit incorporation.

In this study, we applied different genome editing tools to efficiently disrupt repressors BSs in the *HBG1/2* promoters and simultaneously insert a 18-bp long TAL1:GATA1 composite motif, which is recognized by two potent transcriptional activators TAL1 and GATA1^30,31^ to upregulate HbF. We demonstrated that replacing the BCL11A BS with the TAL1:GATA1 motif in SCD HSPCs, increased HbF levels and corrected the sickling phenotype in the erythroid progeny, outperforming approaches that only disrupted the BCL11A BS. Notably, the Cas9-HDR-based strategies yielded a higher frequency of precise edits than prime editing approaches, despite extensive protocol optimizations.

## RESULTS

### PEn outperforms PE to replace repressor BSs with a TAL1:GATA1 motif in the *HBG1/2* promoters

We developed editing strategies that aim at replacing repressors BSs with a composite 18-bp-long TAL1:GATA1 motif recognized by two potent transcriptional activators (**Figure 1A**) to generate a modified γ-globin promoter with increased HbF production. This motif is present in genes highly expressed during erythroid development^30,31^.

**Figure 1:**
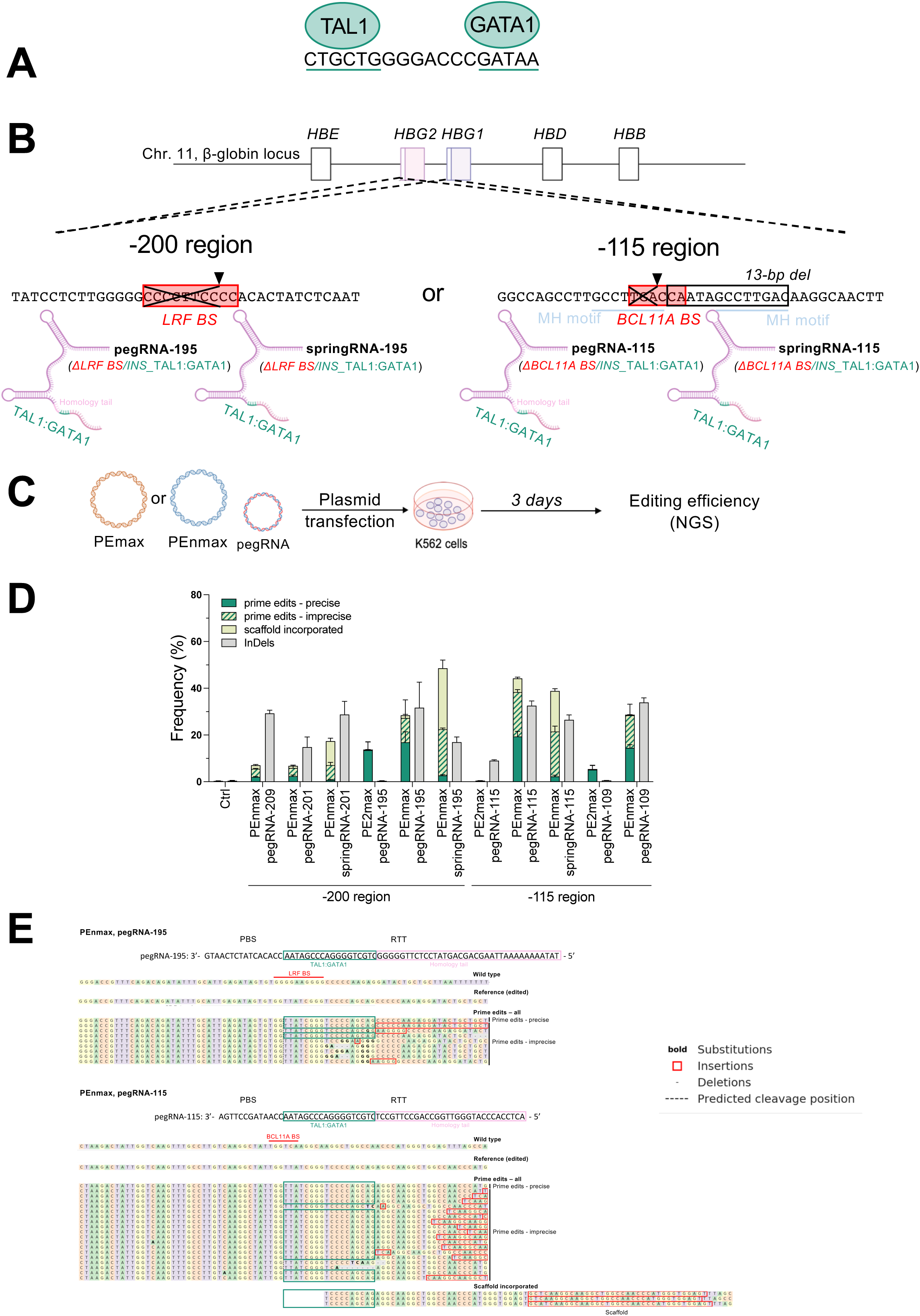
Design and screening of pegRNAs targeting the *HBG1/2* promoters in K562 cells. (A) Representation of the *in vitro* consensus sequence of the composite 18-nt long TAL1:GATA1 motif. (B) Schematic representation of the β-globin locus on chromosome (chr.) 11 including the β-like globin genes (*HBE*, *HBG1*, *HBG2*, *HBD* and *HBB*) and the *HBG1*/*2* promoters. The LRF and BCL11A repressor binding sites (BSs) are highlighted by a light red box. The pegRNA-195 and springRNA-195 and the pegRNA-115 and springRNA-115, targeting respectively the −200 and the −115 region of the *HBG1/2* promoters, allow the replacement of the LRF or the BCL11A repressor BSs (ΔBCL11A or ΔLRF BS) with the consensus TAL1:GATA1 sequence (INS_TAL1:GATA1). SpringRNAs do not contain the homology tail. The 13-bp HPFH deletion (13-bp del), indicated by an empty black box, occurs via alt-EJ after DSB because of the 8-bp micro-homology motifs (MH motifs, blue lines) located in the −115 region^23^. Black arrowheads indicate the corresponding springRNA/pegRNA’s cleavage sites at position −195 and −115 from the transcriptional start site (TSS) of the *HBG1/2* genes. (C) Experimental protocol used to test prime editing strategies in K562 cells. Plasmids expressing PEnmax or PE2max and a pegRNA or a springRNA were co-transfected in K562 cells. The prime editing efficiency is assessed by PCR amplification of the *HBG1/2* promoters and NGS 3 days after transfection. Data were analyzed with the CRISPResso2 webtool. Samples transfected with TE (Tris-EDTA) buffer were used as controls (Ctrl). (D) Percentage of NGS reads containing the prime editing events (precise, imprecise, scaffold incorporated) and InDels. Bars represent the mean ± SD of 3 biological replicates. (E) CRISPResso2 alignments of the *HBG1/2* promoter sequences, as determined by NGS, in samples edited with pegRNA-195 (upper panel) or pegRNA-115 (lower panel) in combination with PEnmax. We reported the PBS and RTT sequences of the pegRNAs, the wild type sequence (no prime editing event), the reference (i.e., the expected edited sequence used by CRISPResso2 to align NGS reads), all prime edits (including precise and imprecise events) and “scaffold incorporated” editing events (identified for pegRNA-115). The TAL1:GATA1 motif and the homology tail are highlighted with a green and a pink box, respectively. BCL11A and LRF BSs are indicated with a red box in the wild type sequence.

First, we designed several pegRNAs (targeting the LRF and the BCL11A BS in the −200 and −115 regions of the *HBG1/2* promoters, respectively) with spacer sequences used in previously developed sgRNAs, and pegRNAs showing high editing efficiency and low off-target activity in combination with Cas9 nuclease^23,34^ (**Figure 1B**). All pegRNAs carry a RTT (with a length ranging from 48- to 55-nt) containing a total or partial deletion of the BS depending on the sgRNA spacer (inducing a cleavage site inside or outside the repressor BS, respectively), the 18-bp long TAL1:GATA1 sequence, and a 30-nt long homology tail (or 36-nt long for the pegRNA-195) (**Figure 1A** and **1B** and **TableS1**). For pegRNA-201, pegRNA-195, and pegRNA-115, which induce a cleavage site inside the repressor BS, we also designed springRNA to exploit the Primed INSertions strategy (PRINS), which is known to promote insertions at the DSB through NHEJ^35,36^. The springRNAs contain the same spacer and PBS of the corresponding pegRNAs with an RTT carrying the 18-nt TAL1:GATA1 sequence that after reverse transcription is directly ligated at the DSB site by NHEJ (**Figure 1B**). This allows the insertion of the desired motif into the repressor BS, causing its inactivation. We tested the different constructs by plasmid transfection in K562 cells comparing PE2max to the PEnmax (**Figures 1C**). We initially evaluated the editing efficiency by Sanger sequencing. Overall, PEnmax outperformed PE2max in editing both −115 and −200 sites (**Figure S1A**). As expected, PEn induced a higher frequency of InDels compared to PE2max (**Figure S1A**). In addition, none of the three springRNAs outperformed PE2max or PEnmax at inserting the expected motif (**Figure S1A**).

To better characterize the DNA repair profile induced by the most efficient editing strategies, *HBG1/2* promoters were amplified by PCR and subjected to NGS (**Figure 1D**). The frequency of total prime editing events (i.e., precise, imprecise and edits incorporating the scaffold) approached 30% for pegRNA-195 and pegRNA-109, and 45% for pegRNA-115 using the PEnmax, of which the precise edits represent around 17%, 15%, and 19% of total NGS reads, respectively (**Figures 1D)**. As shown by Sanger sequencing, the PEnmax outperformed the PEmax in generating precise and imprecise edits (**Figures 1D**). PRINS (using springRNA and the PEnmax) generated mainly imprecise or edits containing the pegRNA scaffold, confirming that NHEJ facilitates the insertion of long DNA sequences (i.e., including the scaffold) at these target sites (**Figures 1D**). Amongst the three best-performing pegRNAs, we selected pegRNA-195 and pegRNA-115 for further analyses, as these pegRNAs generate a cleavage site inside the repressor BS, favoring its disruption, and are associated with the highest frequency of precise edits. The “imprecise” prime edits observed using pegRNA-195 and pegRNA-115 correspond mainly to the deletion of the repressor BSs and the introduction of a complete or partial TAL1:GATA1 motif with other substitutions, insertions (i.e., duplications of part of the RTT) or deletions (**Figures 1D** and **1E**). The “scaffold incorporated” edits are characterized by integration of the TAL1:GATA1 motif, introduction of the RTT and partial incorporation of the scaffold sequence; these modifications lead also the disruption of the repressor BSs (**Figure 1E**). Consequently, both imprecise and scaffold-incorporated edits are also expected to reactivate γ-globin expression similar to InDels that inactivate the repressor BSs^23^. Furthermore, we observed that the PEn system generates a higher proportion of *HBG1/2* 4.9-kb deletions compared to PEmax-based strategies. This deletion event is the consequence of concurrent cleavage of the two identical *HBG1/2* promoter regions resulting in the loss of the *HBG2* gene, a phenomenon also observed in Cas9 nuclease- and base editor-mediated approaches^23,32^ (**Figure S1B**).

### The prime editing efficiency is modest in primary SCD HSPCs despite the optimization of pegRNA and delivery method

Prior to evaluating PEn strategies in patients’ cells, we further optimized the pegRNA-195 and the pegRNA-115 to facilitate their synthesis– specifically aligning with current protocols optimized for RNA with a maximum length of 150 nt- and to improve the editing efficiency. To meet this length criteria, we generated pegRNA-195_S and pegRNA-115_S, harboring a shorter (S) homology tail of 16- and 7-nt, respectively, and we compared them with the original design constructs (L; **Figure 2A**). We tested pegRNA-195_S and pegRNA-115_S with PEnmax in K562 cells using different delivery methods (i.e., ribonucleoprotein, RNP, or RNA transfection) that are suitable for clinical applications^32,37^. The editing efficiency was evaluated by NGS to quantify the frequency of desired insertion and alternative modifications (**Figure 2A**). The pegRNAs with the shortest homology tail inserted both precise and imprecise edits at higher efficiency compared to long pegRNAs, while the incorporation of the scaffold remained low (**Figures 2B** and **2C**). Overall, the frequency of precise and imprecise edits was greater upon RNA transfection compared to RNP delivery (**Figures 2B** and **2C**). Importantly, the proportion of InDels (likely less productive in HbF reactivation compared to precise and imprecise edits) was reduced using short pegRNAs upon RNA delivery (**Figures 2B** and **2C**). The frequency of 4.9-kb deletion, measured by ddPCR, was comparable between S and L constructs and higher in RNA-transfected samples (**Figures 2B** and **2C**). These results suggest that RNA delivery, by increasing pegRNA/PEn concentration in the cells compared to RNP transfection, may facilitate the simultaneous cleavage of the two promoters (**Figures 2B** and **2C**).

**Figure 2:**
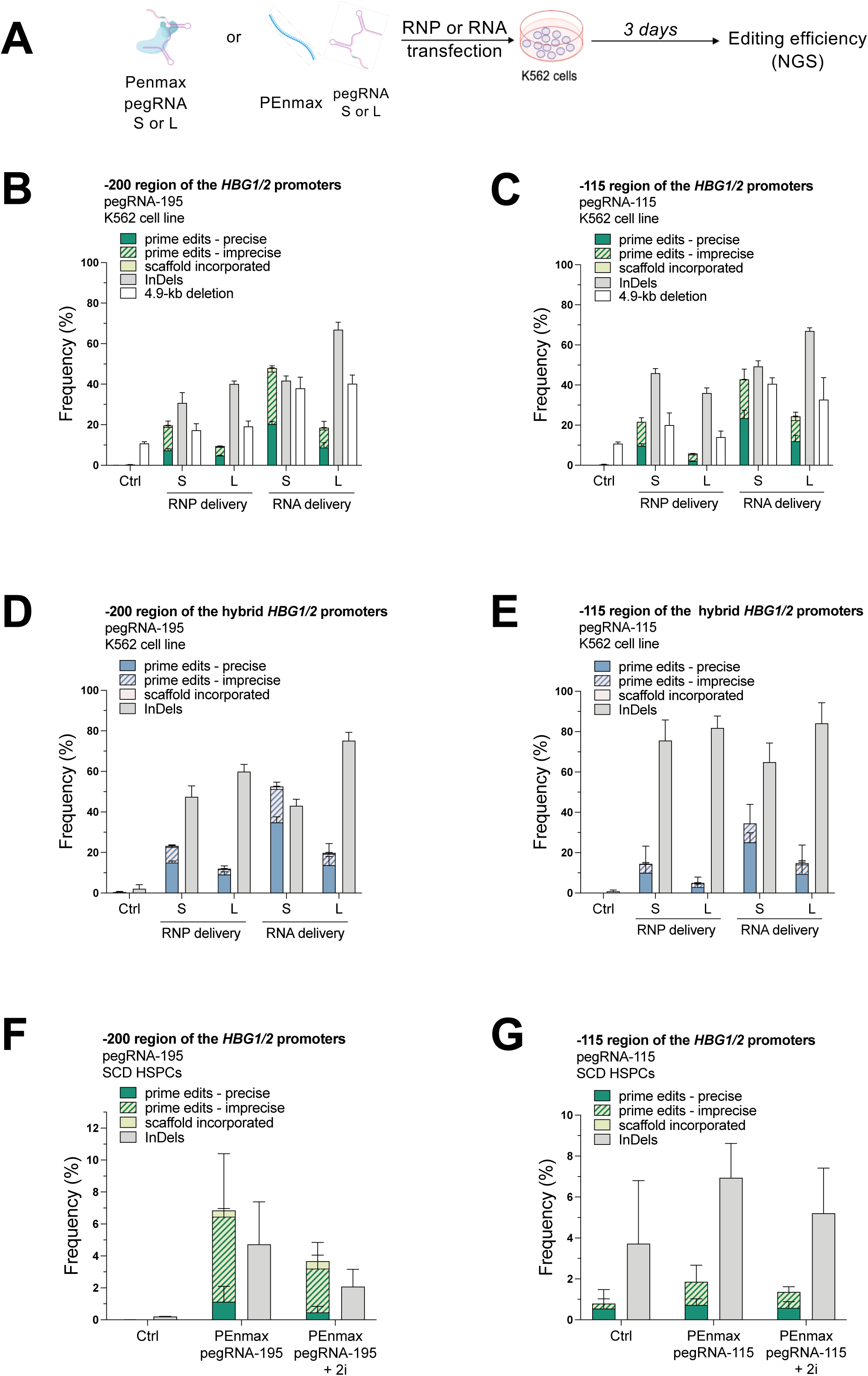
Optimization of the pegRNA constructs and the delivery method in K562 cells. (A) Experimental protocol used for PEnmax experiments in K562 cells. PEnmax and a pegRNA (with a short (S) or a long (L) homology tail) are delivered as RNA or ribonucleoprotein (RNP) in K562 cells. The prime editing efficiency is assessed by NGS 3 days after transfection and analyzed by CRISPResso2. Samples transfected with TE (Tris-EDTA) buffer were used as controls (Ctrl). (B, C) Percentage of the total *HBG1/2* promoters containing the total prime edits (precise, imprecise, scaffold incorporated) and InDels induced by (B) pegRNA-195_S or pegRNA-115_L, or (C) pegRNA-115_S or pegRNA-115_L, delivered as RNA or RNP in K562 cells. We report also the frequency of the 4.9-kb deletion measured by ddPCR for control (Ctrl, TE-transfected cells) and prime-edited conditions. Bars represent the mean ± SD of 3 biological replicates. (D, E) Percentage of the hybrid *HBG1/2* promoters containing the prime edits (precise, imprecise, scaffold incorporated) and InDels induced by (D) pegRNA-195_S or pegRNA-115_L or (E) pegRNA-115_S or pegRNA-115_L, delivered as RNA or RNP in K562 cells. Bars represent the mean ± SD of 3 biological replicates. (F) Percentage of total *HBG1/2* promoters containing the total prime edits (precise, imprecise, scaffold incorporated) and InDels induced by (left) pegRNA-195_S or (right) pegRNA-115, delivered as RNA in HSPCs from patients with SCD treated or not with DNA repair inhibitors (2i). After electroporation, SCD HSPCs were cultured for 6 days in pre-activation medium before sequencing. The *HBG1/2* promoters were amplified and subjected to NGS. Data were analyzed with the CRISPResso2 webtool. Bars represent the mean ± SD of 3 replicates of non-mobilized SCD CD34^+^ cells from 2 different SCD donors.

Subsequently, we characterized the editing profile of the hybrid *HBG1/2* promoters - those generated following the 4.9-kb deletion by NGS to evaluate whether these novel promoters can induce robust γ-globin expression (**Figures 2D** and **2E**). For pegRNA-195 the total frequency of precise and imprecise events was similar to that observed in the total *HBG1/2* promoter population, while pegRNA-115 showed a slightly lower overall editing frequency. Notably, the relative proportion of precise versus imprecise edits was increased in the hybrid promoters compared to the total promoters (**Figures 2B, 2C, 2D** and **2E**). In most cases, the incidence of InDels was higher in hybrid *HBG1/2* promoters (compared to total promoters), except for pegRNA-195_S in RNA-transfected cells, which exhibited the highest frequencies of both precise and imprecise edits (**Figures 2D** and **E**). These results indicate that, despite the generation of *HBG1/2* 4.9-kb deletions using PEn based strategy, the resulting hybrid promoters remain capable of activating γ-globin expression either by installation of activating motifs or by generation of InDels that disrupt the repressor BSs (**Figures 2D** and **E**).

To favor the incorporation of precise edits and maximize γ-globin reactivation, we tested the treatment with DNA-PKc inhibitor (DNA-PKi, AZD7648) and Polθ inhibitor (Polθi, ART558) that inhibit NHEJ and alt-EJ, respectively (**Figures S1C, S1D, S1E** and **S1F**). The DNA-PKi was tested alone and in combination with Polθi in K562 cells (2i). The combined inhibition increased product purity and diminished the frequency of InDels, with the best-performing conditions being those using short pegRNA and RNA delivery (**Figures S1C** and **D**). Similar results were observed by NGS of the hybrid *HBG1/2* promoters (**Figures S1E** and **F)**.

Finally, we tested our best-performing strategies (i.e., pegRNA-195_S and pegRNA-115_S) by RNA transfection in HSPCs obtained from patients with SCD. Editing efficiencies were low for the two strategies (with or without DNA-PKc and Polθ inhibitors) reaching only 6.4% and 2.9% of total prime edits for pegRNA-195_S and pegRNA-115_S, respectively (1.1% of precise edits; **Figures 2F** and **2G**). In addition, the proportion of InDels approached or exceeded the frequency of prime editing events (**Figures 2F** and **2G**).

### Cas9-HDR outperforms PEn in replacing the BCL11A BS by the TAL1:GATA1 motif, offering enhanced precision and reduced InDels via DNA repair modulation

To increase the generation of precise edits, we developed a CRISPR/Cas9 HDR strategy (referred as Cas9-HDR) to replace the BCL11A repressor BS with the TAL1:GATA1 motif in the −115 region of the *HBG1/2* promoters. We used the sgRNA-115 that has been previously shown to disrupt the BCL11A BS using Cas9 nuclease and reactivate HbF^23^. We designed several ssODN templates containing the partial BCL11A BS deletion, the TAL1:GATA1 sequence (**Figure 1A**), and homology arms (i.e., ranging from 40 to 110 nt on each extremity of the ssODN) complementary to the (+) or (-) DNA strand with the target site being localized on the (+) strand (**Figures 3A** and **3B**).

**Figure 3:**
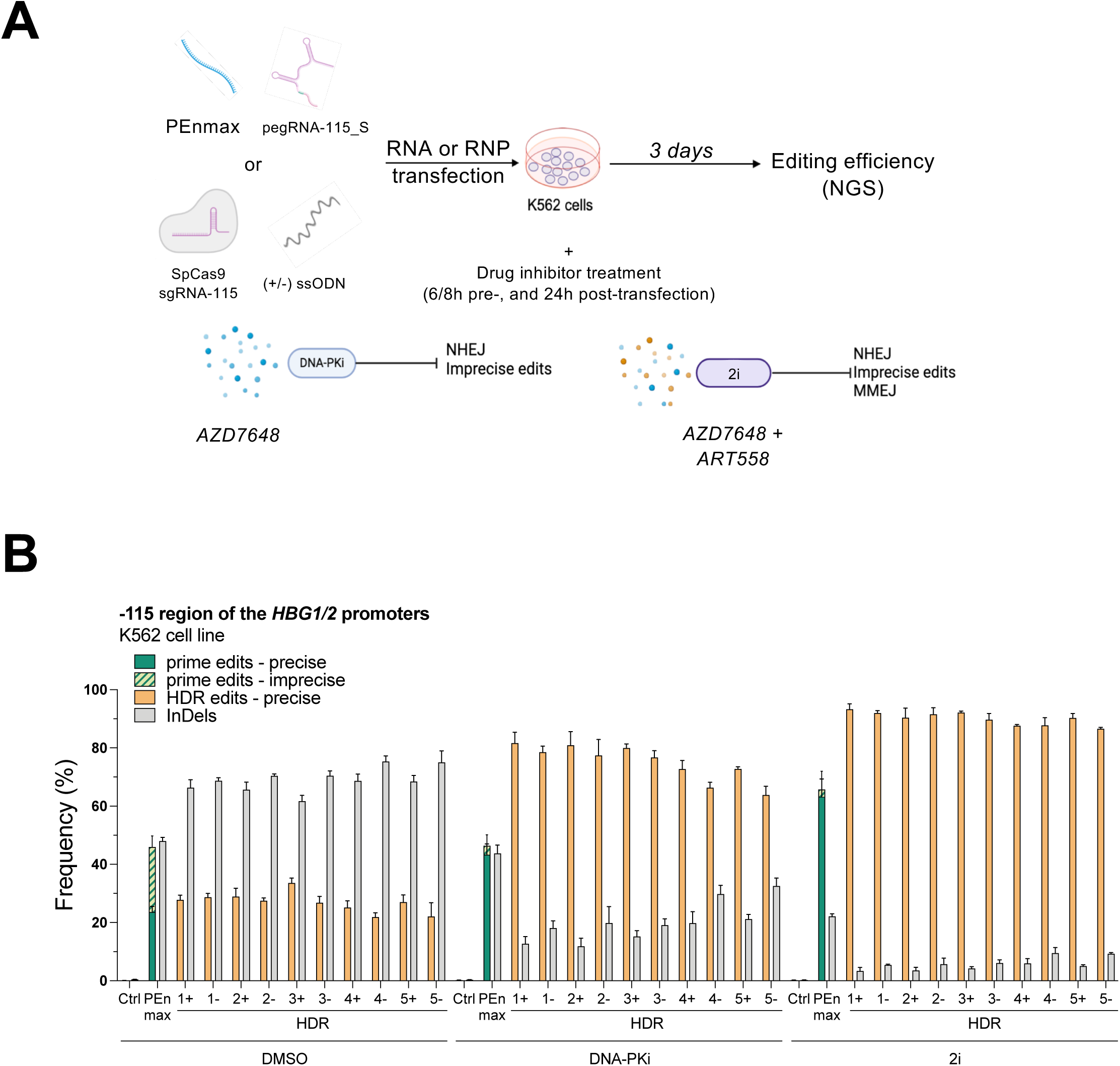
Design and screening of ssODNs to replace the BCL11A BS with the TAL1:GATA1 motif using a Cas9-HDR strategy in K562 cells. (A) Experimental protocol used for PEnmax and Cas9-HDR experiments in K562 cells. PEnmax and pegRNA-115_S were delivered as RNA and Cas9/sgRNA as RNP in K562 cells. Cells were treated with DMSO, 1 μM of AZD7648 (DNA-PKi), or with 1 μM of AZD7648 and 3 μM of ART558 (2i). The editing efficiency was assessed by NGS 3 days after transfection. Samples transfected with TE (Tris-EDTA) buffer were used as controls (Ctrl). (B) Percentage of total *HBG1/2* promoters containing the total prime edits (precise, imprecise), the HDR edits (precise), and InDels induced by pegRNA-115_S and PEnmax, or sgRNA-115, Cas9 and ssODNs, in the presence or absence of different inhibitors. Bars represent the mean ± SD of 3 biological replicates.

First, we screened the different ssODNs in K562 cells by RNP transfection, a clinically approved method to deliver CRISPR/Cas9 reagents in HSPCs^38,39^ (**Figure 3A**). In parallel, we compared the PEn/pegRNA115_S strategy using the best delivery method of the editing tools, i.e. RNA delivery (**Figure 2D**). Virtually all ssODNs generated the precise edits at a slightly higher level compared to the PEn strategy (**Figure 3B**). The Cas9-HDR strategies showed greater InDel frequencies in comparison to the PEn strategy (**Figure 3B**). To maximize γ-globin reactivation by reducing InDels and favoring the incorporation of precise edits we tested DNA-PKi (AZD7648) and Polθi (ART558)^24,40^. The DNA-PKi led to greater increase of precise edits and reduction of InDels in cells treated with Cas9-HDR compared to PEn-treated samples (**Figure 3B**). The combination of the two inhibitors (2i) further increased the product purity (i.e., the proportion of precise edits over InDels) for both Cas9-HDR and PEn strategies (**Figure 3B**). The inhibition of the alt-EJ pathway is of particular importance in the −115 region that contains 8-bp MH motifs at the target site that may induce alt-EJ after DSB, thus generating at high frequency a 13-bp deletion, as we previously demonstrated^23^ (**Figure 1A**). As an example, cells edited with Cas9, sgRNA-115 and ssODN (1+) showed ∼6%, ∼8%, and ∼1% of 13-bp deletion without inhibitors, with DNA-PKi alone, and with DNA-PKi in combination with Polθi, respectively. A similar trend was observed in PEn/pegRNA115_S-treated cells (∼9%, ∼31%, and ∼12%, without inhibitors, with DNA-PKi alone, and with DNA-PKi+Polθi, respectively). Amongst all ssODNs, we selected the ssODN (1+) that led to the highest editing efficiency (93.3%) and the lower InDel frequency (3.4%) after 2i treatment in K562 cells (**Figure 3B**).

### The Cas9-HDR strategy efficiently reactivates γ-globin expression and corrects the sickling phenotype in RBCs differentiated from SCD HSPCs

We tested the Cas9-HDR strategy in primary cells derived from patients with SCD to evaluate if this combined approach (i.e., disruption of a repressor BS and insertion of two activator BSs) reactivates HbF expression to a higher level compared the sole disruption of the repressor BS^23,41,42^. HSPCs were transfected with RNPs containing sgRNA-115 and Cas9, with or without (1+) ssODN (**Figure 4A**). In addition, for the Cas9-HDR strategy, the cells were incubated with or without 2i for 24 h to promote precise edits and reduce InDels, as shown in K562 cells (**Figures 3B** and **4A**). In the absence of the ssODN, the DSB is mainly repaired by NHEJ and alt-EJ leading to the formation of InDels (67.9%) and the disruption of the BCL11A BS, as previously demonstrated^23,41,42^ (**Figure 4B**). In liquid erythroid cultures, we generated 9.8% and 17.0% of precise edits in Cas9-HDR-treated cells without and with 2i, respectively, exceeding the frequency obtained using the PEn strategy (**Figures 2F** and **4B**). In addition, InDel frequency was significantly reduced upon 2i treatment (from 38.8% to 16.9%; **Figure 4B**). The frequency of *HBG1/2* 4.9-kb deletion was similar in all the samples (**Figure 4B**). Finally, we assessed the off-target activity by sequencing the major off-target site of sgRNA-115 (mapping to an intergenic region on chromosome 2) that we previously identified using GUIDE-seq^23^. Importantly, NGS did not reveal any InDels in Cas9-, Cas9-HDR, or 2i-Cas9-HDR-treated conditions (**Figure 4C**).

**Figure 4:**
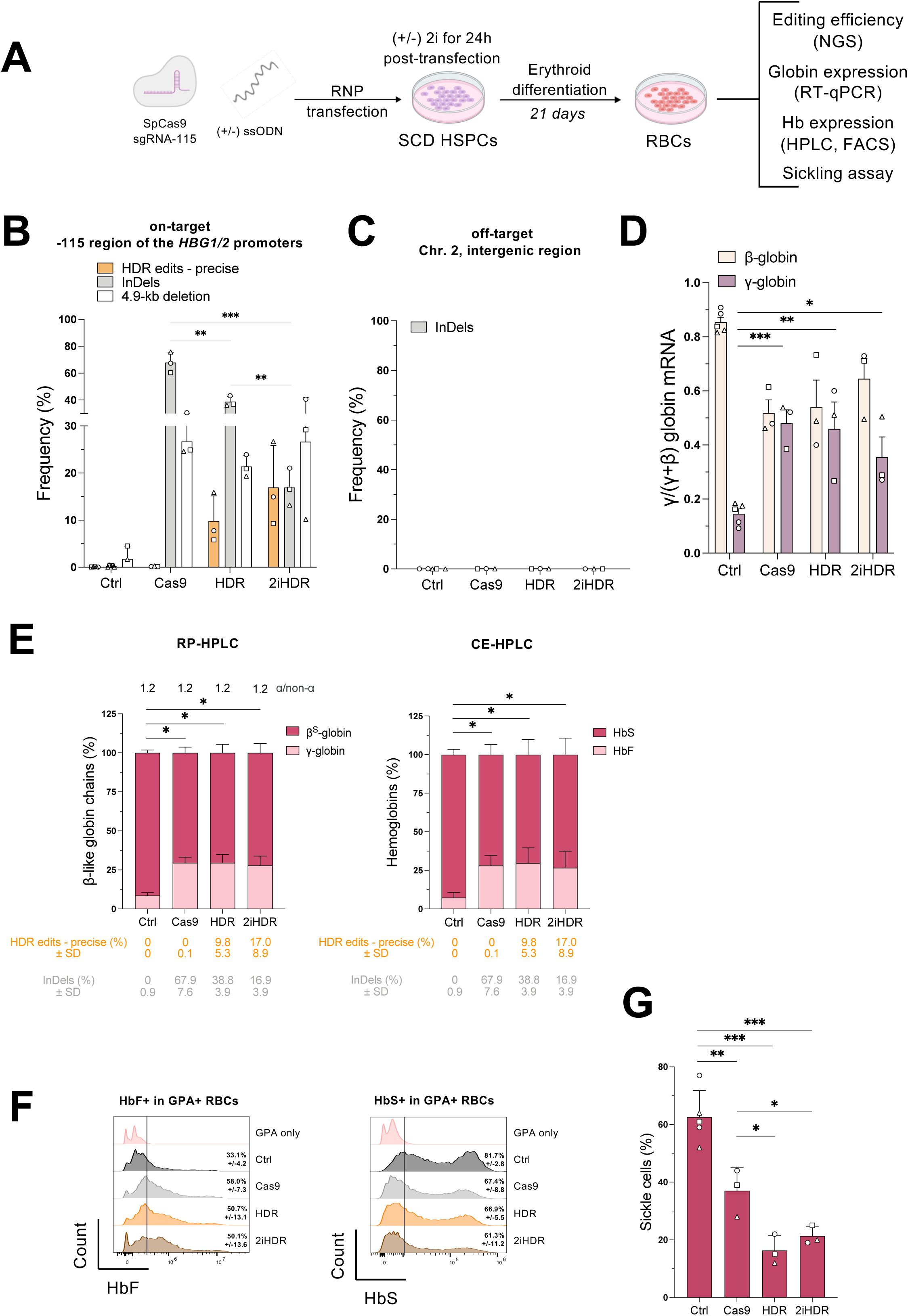
The replacement of the BCL11A BS by the TAL1:GATA1 motif reactivates HbF expression and corrects the sickling phenotype in RBCs differentiated from SCD HSPCs. (A) Experimental protocol used for the Cas9-based strategies in HSPCs derived from patients with SCD. Cas9 and sgRNA-115 were delivered as RNP complexes with or without ssODN (1+) in non-mobilized peripheral blood SCD HSPCs (n=3 independent biological experiments; 2 different donors). After transfection, cells were treated for 24 h with 2i (2iHDR) or DMSO (as negative control), then differentiated for 21 days in mature RBCs and subjected to different assays. (B) Percentage of total *HBG1/2* promoters containing the HDR edits (precise) and InDels, induced by sgRNA-115 and Cas9 +/- ssODN (1+) and 2i. The frequency of the 4.9-kb deletion was measured by ddPCR. Samples transfected with TE (Tris-EDTA) buffer or transfected with the Cas9 protein only were used as controls (Ctrl). Bars represent the mean ± SD of 3 independent biological experiments (2 different donors). Statistical significance was assessed using unpaired t-test. ***p<0.001, **p<0,005. (C) Editing at the main off-target site (chromosome 2, intergenic region), identified by GUIDE-seq^23^ in control (Ctrl, TE- and transfected with the Cas9 protein), Cas9- and HDR-treated conditions. Bars represent the mean ± SD of 3 independent biological experiments (2 different donors). (D) RT-qPCR analysis of β-like globin mRNA levels in patients’ erythroblasts at day 13 of erythroid differentiation. β-like globin mRNA expression was normalized to α-globin mRNA. Bars represent the mean ± SD of 3 independent biological experiments (2 different donors). Statistical significance was assessed using unpaired t-test. ***p<0.001, **p<0.005, *p<0.05. (E) Expression of (left) β^S^- and γ-globin chains, and (right) HbS and HbF measured by RP-HPLC and CE-HPLC, respectively, in RBCs differentiated from SCD HSPCs. We calculated the proportion of each β-like globin over the total β-like globin chains. Bars represent the mean ± SD of 3 independent biological experiments from 2 different donors. The ratio α/non-α globin is reported on top and HDR precise edits and InDels are reported on the bottom. Statistical significance was assessed using Mann-Whitney test. *p<0.05. (F) Flow cytometry histograms showing the percentage of (left) HbF- and (right) HbS-expressing cells in the GPA^+^ population for samples stained only with the GPA antibody (GPA only) and for control (TE-transfected) and edited samples. Frequency ± SD is indicated for 3 independent biological experiments from 2 different donors. (G) Frequency of sickled RBCs upon O_2_ deprivation in control (Ctrl, TE- and transfected with the Cas9 protein) and edited samples. Cells were counted by a blinded observer for all conditions (>300 total randomly selected cells per condition). Bars represent the mean ± SD of 3 independent biological experiments (2 different donors). Statistical significance was assessed using unpaired t-test. ***p<0.001, **p<0.005, *p<0.05.

Next, we differentiated control and treated cells in mature RBCs. The enucleation rate as well as the expression of erythroid markers were similar between control and edited samples, demonstrating no impact on the erythroid differentiation (**Figure S2A-C**). RT-qPCR, RP-HPLC, and CE-HPLC analyses showed a strong and significant γ-globin reactivation in all edited samples, at both RNA and protein levels (**Figure 4D-E**). Of note, the ratio between α and non-α globin chains was not altered in any of the samples (**Figure 4E**). Although the overall editing efficiency was lower in Cas9-HDR or 2i-Cas9-HDR-treated cells compared to Cas9 samples, we observed similarly high γ-globin and HbF levels across all three conditions (**Figure 4E**). Flow cytometry analysis also showed an increased frequency of HbF+ cells and a decrease in the fraction of HbS-expressing cells in all edited samples (**Figure 4F**). These results suggest that precise edits generated by the Cas9-HDR approach are associated with great γ-globin reactivation (**Figure 4E**).

Finally, we incubated mature RBCs under hypoxic conditions to induce sickling (**Figure 4G**). Cas9 treatment led to a 1.7-fold decrease in the frequency of sickle cells, and the Cas9-HDR approaches further reduced RBC sickling (**Figure 4G**).

### A clonal analysis confirmed safety and efficacy of the HDR-based strategy replacing the BCL11A BS with the TAL1:GATA1 activating motif

HSPCs from patients with SCD were transfected with and without ssODN and 2i and plated in a semi-solid medium allowing erythroid (BFU-E) and granulocyte/monocyte (CFU-GM) differentiation at clonal level (CFC assay; **Figure 5A**). Control and edited samples harbored a similar clonogenicity potential, although 2i led to a reduction of erythroid colonies for 2 out of 3 donors (**Figure 5B**). Editing efficiency in BFU-E and CFU-GM pools were similar to that obtained in liquid erythroid cultures (**Figures 4B** and **5C-D**).

**Figure 5:**
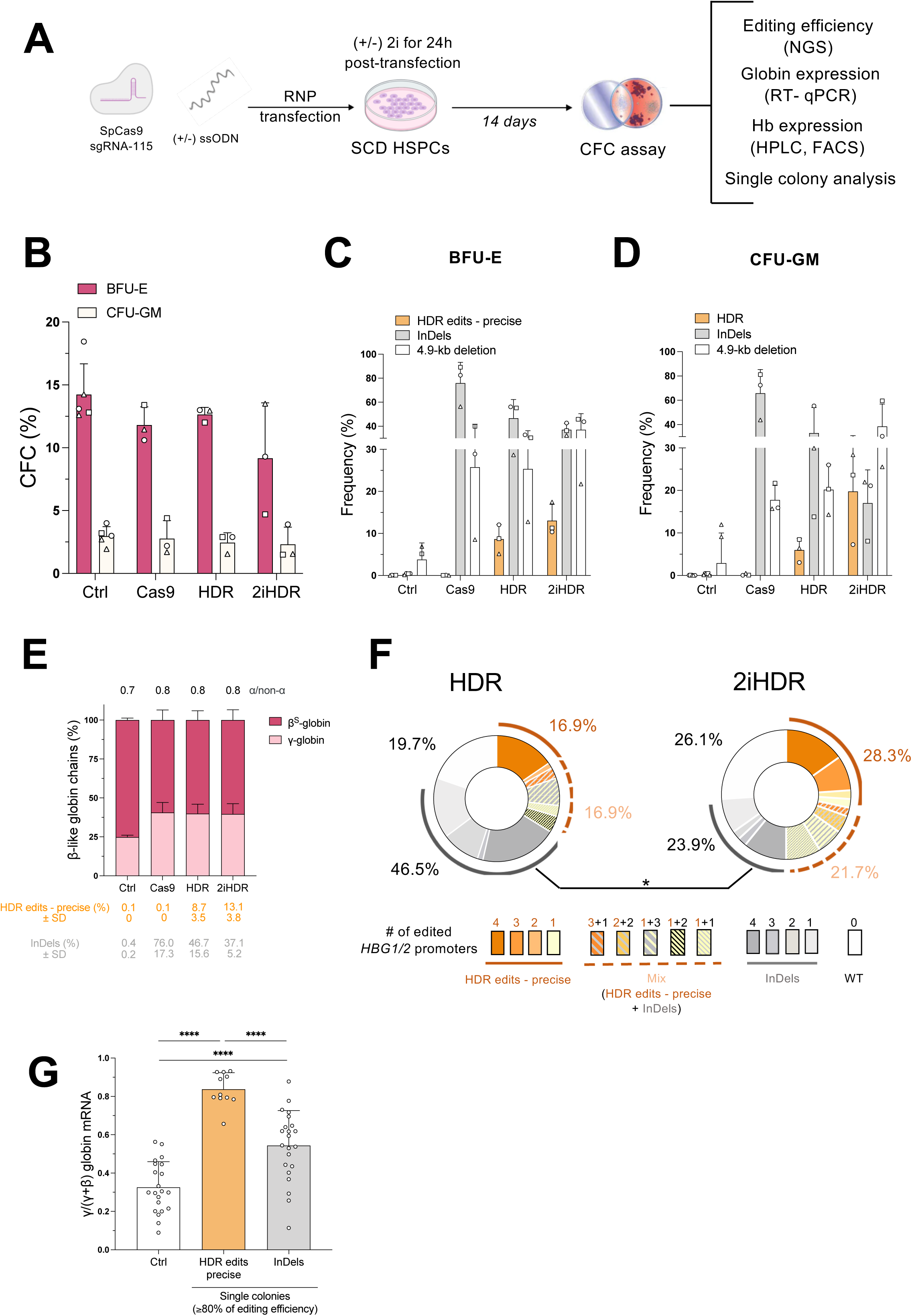
Viability, editing, and *HBG1/2* expression in progenitors derived from SCD HSPCs. (A) Experimental protocol used for the Cas9-based strategies in HSPCs derived from patients with SCD. Cas9 and sgRNA-115 were delivered as RNP complexes with or without ssODN (1+) in non-mobilized peripheral blood SCD HSPCs (n=3 independent biological experiments; 2 different donors). After transfection, cells were treated for 24 h with 2i (2iHDR) or DMSO (as negative control), and then plated in a semi-solid medium allowing the growth of BFU-E and CFU-GM for 14 days. (B) BFU-E and CFU-GM frequencies for control (Ctrl, TE- and transfected with the Cas9 protein only), Cas9- and Cas9-HDR-treated conditions. Bars represent the mean ± SD of 3 independent biological experiments (2 different donors). No statistical differences were observed between edited samples using Mann-Whitney. (C, D) Percentage of total *HBG1/2* promoters in (C) BFU-E or (D) CFU-GM colonies, containing the HDR edits (precise), or InDels in Ctrl (TE- and transfected with the Cas9 protein only), Cas9- and Cas9-HDR-edited conditions +/- the 2i, as evaluated by NGS. The frequency of the 4.9-kb deletions was measured by ddPCR. Bars represent the mean ± SD (n=3 independent biological experiments; 2 different donors). No statistical differences were observed between edited samples using Mann-Whitney. (E) Expression of β- and γ-globin chains measured by RP-HPLC in BFU-E pools derived from SCD HSPCs. We calculated the proportion of each β-like globin over the total β-like globin chains. Bars represent the mean ± SD of 3 independent biological experiments (2 different donors). The ratio α/non-α globin is reported on top of the graph and HDR precise edits and InDels are reported below the graph. No statistical differences were observed between edited samples using Mann-Whitney. (F) Frequency of BFU-E colonies carrying precise HDR edits, precise HDR edits and Indels, Indels, or WT promoters in HDR- or 2iHDR-treated conditions [n=3 independent biological experiments (2 different donors); n= 71 and 46 single colonies, respectively]. Edited colonies contain 1 to 4 γ-globin promoters either with only the HDR edits (precise, orange), only InDels disrupting the BCL11A BS (grey), or a mix of all of HDR edits and InDels (orange + grey). Statistical significance was assessed using two-way ANOVA with Sidak’s multiple comparison test. *p<0.05. (G) RT-qPCR analysis of γ-globin expression in single BFU-E derived from control (Ctrl, TE- and transfected with the Cas9 protein only) or edited cells. Edited colonies with ≥80% of editing efficiency were selected. The γ-globin mRNA expression was normalized to α-globin. Bars represent the mean ± SD of 11 to 22 single BFU-E from 3 independent biological experiments (2 different donors). Statistical significance was assessed using unpaired t-test. ****p<0.0001.

Edited BFU-E bulk populations showed γ-globin reactivation (**Figures 5E**). To better characterize the proportion of precise edits and InDels, we evaluated the genotype of single BFU-Es derived from Cas9-HDR- and 2i-Cas9-HDR-edited HSPCs (**Figure 5F**). We observed that 2i significantly reduced the proportion of BFU-Es carrying alleles containing InDels (23.9%) compared to untreated cells (46.5%; **Figure 5F**). In addition, 2i favored the incorporation of the desired edits, leading to a higher proportion of clones harboring alleles with the precise edits in 1 to 4 *HBG1/2* promoters and clones with both precise edits and InDels (**Figure 5F**).

To better evaluate the difference in γ-globin output associated with the combined HDR strategy and the sole Cas9-mediated disruption of the BCL11A BS, we measured γ-globin expression at mRNA level in single BFU-E clones harboring only the precise edits or InDels (**Figures S3A** and **5G**). We observed a positive correlation between the frequency of the precise edits and γ-globin reactivation (R^2^ = 0.68), while for InDel-carrying clones, γ-globin levels were more heterogenous (R^2^ = 0.18, **Figure S3A**). We then compared γ-globin expression in erythroid clones with a similar editing efficiency (≥80%, i.e., mainly clones with 4 edited promoters). Importantly, γ-globin reactivation was significantly higher in clones carrying the precise edits, i.e. the TAL1:GATA1 motif and the disruption of the BCL11A BS, than in clones harboring only InDels disrupting the BCL11A repressor BS (**Figure 5G**).

## DISCUSSION

Recent studies have demonstrated great potential in achieving high levels of therapeutic γ-globin expression to treat β-hemoglobinopathies. The exagamglogene autotemcel (known as Casgevy) is the first FDA-approved CRISPR-intervention that disrupts the *BCL11A* erythroid enhancer to treat patients with β-hemoglobinopathies^38,39^. However, the clinical study targeting the *BCL11A* enhancer showed variability in the extent of HbF reactivation among individuals, relatively low levels of Hb with HbS still accounting for a large proportion of the total Hb and only modest correction of ineffective erythropoiesis^38,39^, the latter potentially due to the BCL11A role in erythroid development^16,17^. Recent work based on the disruption of the BCL11A repressor BS in the −115 region in the *HBG* promoters^20–22^ showed significant increase in HbF levels (comparable to the Casgevy approach) with no detectable off-target activity^43–45^. This strategy is currently being tested in clinics in patients with β-hemoglobinopathies^44–46^. While the disruption of the BCL11A BS could recapitulate the effect of several benign HPFH mutations, in this study we envisioned a novel strategy to further boost HbF expression by simultaneously inserting a TAL1:GATA1 motif (known to recruit potent activators in erythroid cells) combined to the disruption of the BCL11A repressor BS. This could give a better selective advantage to erythroid cells, thus requiring a lower fraction of edited HSPCs to achieve a therapeutic benefit^47^ (a scenario that is possible with an *in vivo* HSPC-targeting strategy^48^).

We first exploited the PEn system, which allows the incorporation of long DNA stretches in the genome^29^ to replace the LRF or the BCL11A repressor BS, with a 18-bp long TAL1:GATA1 motif present in loci highly transcriptionally active during erythropoiesis^30,31^. The PEn_max_ yielded a higher proportion of total edits, including precise and imprecise edits, compared to the original PE2 system, reaching almost 40% of total edits at the −115 site of the *HBG1/2* promoters. Approximately half of these edits represented precise edits (i.e., precise deletion and insertion), while the remaining edits comprised partial insertion of the TAL1:GATA1 motif, or unintended incorporation of part of the pegRNA RTT/scaffold in the genome. Importantly, imprecise edits are also expected to cause γ-globin reactivation as they disrupt the repressor BS and/or facilitate the recruitment of either TAL1 or GATA1. We believe that PEn outperformed the classical PE system in terms of editing efficiency likely because of the kinetics of the resolution of the single-stranded breaks (SSBs) vs DSBs^49,50^. Specifically, rapid cellular repair of SSB may prevent the incorporation of the reverse transcribed 3’ flap into the genome characteristic of PE strategy, whereas DSB repair leads to slower resolution, favoring the desired insert by the PEn system.

Due to the use of a nuclease, the PEn system led to the formation of InDels that could nevertheless disrupt the repressor BS, mimicking the Cas9-mediated strategies developed for reactivating γ-globin^18,20,21^. The DSBs induced by the PEn system could also lead to the deletion of the 4.9-kb *HBG1-HBG2* intervening genomic region and the subsequent loss of the *HBG2* gene (due to the simultaneous cleavage of the two identical *HBG* promoters) at a high frequency that was, as expected, much higher compared to the original PE system. However, clonal analysis in Cas9-edited cells at the −115 region, showed that the hybrid *HBG* promoters reactivate HbF as long as the BCL11A repressor BS in the −115 region is disrupted at the junction site^20,21^, which was also observed in prime-edited samples.

The modulation of DNA repair pathways using DNA-PKi and/or Polθi increased the product purity and lowered InDels, in line with our recent work^51^. Interestingly, InDel frequency was reduced only by dual inhibition, likely because alt-EJ is upregulated upon NHEJ inhibition. Application of DNA-PKi alone or in combination with Polθi led to an increase of 4.9-kb deletion frequency, probably due to slower kinetics of DSB repair, which increases the possibility of simultaneous DSB formation and excision of intervening sequence ^52^.

The quality of reagents was crucial to achieve high editing efficiency. We observed significantly higher editing efficiency with mRNA-mediated PEn delivery in K562 cells, potentially due to sustained editor production.

This comprehensive optimization enabled the selection of optimal gene editing conditions to be tested in primary HSPCs from patients with SCD. Importantly, we demonstrated for the first time the successful replacement of the repressor BS (i.e., LRF or BCL11A) with an 18-nt long TAL1:GATA1 motif in the *HBG1/2* promoters of primary patient cells using PEn. Nevertheless, editing efficiency remained generally low and variable amongst the different donors, as we recently observed using PE and PEn^34^. Further optimization of the prime editing components and their delivery may be required to achieve clinically relevant editing frequencies^53,54^. Additionally, elucidating mechanisms underlying donor-to-donor variability may inform strategies to achieve more consistent editing outcomes.

We subsequently employed a Cas9-HDR approach to replace the BCL11A BS with the TAL1:GATA1 motif, achieving 90% of precise edits in K562 cells upon 2i treatment compared to 60% achieved with the PEn system. After optimizing protocol parameters in K562 cells, we compared the Cas9-HDR strategy with Cas9-mediated disruption of the BCL11A BS in HSPCs^23,41,42,55^ obtaining up to 20% of targeted insertion events in primary SCD HSPCs with 2i. Importantly, replacing the BCL11A BS with the TAL1:GATA1 activating motif leads to higher and more consistent γ-globin levels than only disrupting the repressor BS (i.e. the therapeutic strategy currently in clinics). These high HbF levels were sufficient to markedly alleviate the sickling phenotype.

No detectable editing was observed at the major predicted off-target site. Cas9-HDR edits did not interfere with erythroid differentiation, although occasional reduction in progenitor numbers was noted. Comprehensive genotoxicity and cytotoxicity assessment are necessary to validate the safety of this strategy. Importantly, we recently demonstrated that co-inhibition of NHEJ and alt-EJ not only enhances editing efficiency and precision, but also reducing off-target editing^51^.

In summary, this study highlights the limitation of prime editing tools for insertion of extended DNA sequences into the genome, particularly in primary cells. Furthermore, our findings establish proof-of-concept that HDR mediated insertion of TAL1:GATA1 motif combined with BCL11A BS disruption yields higher per cell HbF content than current Cas9-based approaches, suggesting a promising approach for the treatment of β-hemoglobinopathies.

## MATERIALS & METHODS

### Cell line culture

Human erythroleukemia K562 cells were maintained at a concentration of 5×10^5^ cells/ml in RPMI 1640 containing glutamine (Gibco) supplemented with 10% fetal bovine serum (Gibco), 2% Hepes (Life Technologies), 1% sodium pyruvate (Life Technologies), and 1% penicillin and streptomycin (Life Technologies) at 37°C and 5%CO_2_.

### CD34^+^ cell purification and culture

We obtained human non-mobilized peripheral blood CD34^+^ HSPCs from SCD patients. SCD samples eligible for research purposes were obtained from the Necker-Enfants malades Hospital (Paris, France). Written informed consent was obtained from all adult subjects. All experiments were performed in accordance with the Declaration of Helsinki. The study was approved by the regional investigational review board (reference: DC-2024-6899, CPP Ile-de-France II “Hôpital Necker-Enfants malades”). HSPCs were purified using the CD34 MicroBead Kit (Miltenyi Biotec). CD34^+^ cells were thawed and cultured for 48h at a concentration of 5×10^5^ cells/ml in the “HSPC medium” containing StemSpan (STEMCELL Technologies) supplemented with penicillin/streptomycin (Gibco), 250 nM StemRegenin1 (STEMCELL Technologies), and the following recombinant human cytokines (PeproTech): human stem-cell factor (SCF) (300 ng/ml), FMS-like tyrosine kinase 3 ligand (Flt-3L) (300 ng/ml), thrombopoietin (TPO) (100 ng/ml), and interleukin-3 (IL-3) (60 ng/ml).

### Plasmids

Plasmids used in this study include:

pMJ920 Cas9-GFP-expressing plasmid (Addgene #42234)

pCMV-PEmax (Addgene #174820)

pCMV-PEnmax (provided by AstraZeneca, Sweden) was generated by gene synthesis (Genscript). The PEnmax corresponds to PEmax with the H840A mutation reversed to the original histidine, restoring nuclease activity.

pU6-pegRNA-GG-acceptor (Addgene #132777)

The PEmax-SpRY and the PEnmax-SpRY plasmids were created by replacing the sequence encoding the PAM-interacting motif (PIM) of the Cas9 with the PIM-encoding sequence of the Cas9-SpRY:

- PIM sequence of the PEmax 5’GTGCAGACAGGCGGCTTCAGCAAAGAGTCTATCCTGCCCAAGAGGAACAGCGATAAGCTGATCGCCAGAAAGAAGGACTGGGACCCTAAGAAGTACGGCGGCTTCGACAGCCCCACCGTGGCCTATTCTGTGCTGGTGGTGGCCAAAGTGGAAAAGGGCAAGTCCAAGAAACTGAAGAGTGTGAAAGAGCTGCTGGGGATCACCATCATGGAAAGAAGCAGCTTCGAGAAGAATCCCATCGACTTTCTGGAAGCCAAGGGCTACAAAGAAGTGAAAAAGGACCTGATCATCAAGCTGCCTAAGTACTCCCTGTTCGAGCTGGAAAACGGCCGGAAGAGAATGCTGGCCTCTGCCGGCGAACTGCAGAAGGGAAACGAACTGGCCCTGCCCTCCAAATATGTGAACTTCCTGTACCTGGCCAGCCACTATGAGAAGCTGAAGGGCTCCCCCGAGGATAATGAGCAGAAACAGCTGTTTGTGGAACAGCACAAGCACTACCTGGACGAGATCATCGAGCAGATCAGCGAGTTCTCCAAGAGAGTGATCCTGGCCGACGCTAATCTGGACAAAGTGCTGTCCGCCTACAACAAGCACCGGGATAAGCCCATCAGAGAGCAGGCCGAGAATATCATCCACCTGTTTACCCTGACCAATCTGGGAGCCCCTGCCGCCTTCAAGTACTTTGACACCACCATCGACCGGAAGAGGTACACCAGCACCAAAGAGGTGCTGGACGCCACCCTGATCCACCAGAGCATCACCGGCCTGTACGAGACACGGATCGACCTGTCTCAGCTGGGAGGTGAC3’
- PIM sequence of the PEmax-SpRY 5’GTGCAGACAGGCGGCTTCAGCAAAGAGTCTATCAGACCCAAGAGGAACAGCGATAAGCTGATCGCCAGAAAGAAGGACTGGGACCCTAAGAAGTACGGCGGCTTCCTGTGGCCCACCGTGGCCTATTCTGTGCTGGTGGTGGCCAAAGTGGAAAAGGGCAAGTCCAAGAAACTGAAGAGTGTGAAAGAGCTGCTGGGGATCACCATCATGGAAAGAAGCAGCTTCGAGAAGAATCCCATCGACTTTCTGGAAGCCAAGGGCTACAAAGAAGTGAAAAAGGACCTGATCATCAAGCTGCCTAAGTACTCCCTGTTCGAGCTGGAAAACGGCCGGAAGAGAATGCTGGCCTCTGCCAAGCAGCTGCAGAAGGGAAACGAACTGGCCCTGCCCTCCAAATATGTGAACTTCCTGTACCTGGCCAGCCACTATGAGAAGCTGAAGGGCTCCCCCGAGGATAATGAGCAGAAACAGCTGTTTGTGGAACAGCACAAGCACTACCTGGACGAGATCATCGAGCAGATCAGCGAGTTCTCCAAGAGAGTGATCCTGGCCGACGCTAATCTGGACAAAGTGCTGTCCGCCTACAACAAGCACCGGGATAAGCCCATCAGAGAGCAGGCCGAGAATATCATCCACCTGTTTACCCTGACCAGACTGGGAGCCCCTAGAGCCTTCAAGTACTTTGACACCACCATCGACCCCAAGCAGTACAGAAGCACCAAAGAGGTGCTGGACGCCACCCTGATCCACCAGAGCATCACCGGCCTGTACGAGACACGGATCGACCTGTCTCAGCTGGGAGGTGAC3’

All plasmids were optimized to contain a T7 promoter compatible with the TriLink’s CleanCap-AG Cap1 analogue and a 3’UTR that we previously described^32^.

All pegRNAs were generated by gene synthesis and cloned in pMA vector together with an upstream U6 promoter (GeneArt).

### PEnmax mRNA *in vitro* transcription

Capped PEnmax mRNA was generated following T7 directed *in vitro* transcription using a linearized PEnmax DNA template. The *in vitro* transcription reaction produced mRNA fully modified, replacing uridines with N1-Methyl-pseudouridine, and cap-1 capped using TriLink’s CleanCap-AG Cap1 analogue. The mRNA was subsequently column purified using MEGAClear transcription clean-up kit (ThermoFisher) and mRNA purity was analyzed using a fragment analyzer (Agilent).

### Synthetic pegRNAs, sgRNAs and ssODN constructs

The PBS and RTT were manually designed for all pegRNAs, for which we selected the recommended PBS length of 13-nt, except for pegRNA-195 that was designed using the Prime Design webtool with a 16-nt-long PBS. All pegRNAs carry a RTT (with a length ranging from 48- to 55-nt) containing a total or partial deletion of the LRF or the BCL11A BSs, the 18-bp long TAL1:GATA1 sequence, and a 30-nt long homology tail (or 36-nt long for the pegRNA-195 as defined with the Prime Design webtool). Chemically modified synthetic pegRNAs, sgRNAs and single-stranded oligo-DNA nucleotides (ssODN) were ordered from Integrated DNA Technologies (IDT). Each construct harbored 2’-*O*-methyl analogs and 3’-phosphorothioate non hydrolysable linkages at the first three 5’ and 3’ nucleotides. All pegRNAand and ssODNs sequences used in this study are respectively listed in **Table S1** and **S2**. The target sequence of the sgRNA-115 is 5’-CTTGTCAAGGCTATTGGTCA**AAG**-3’, with the protospacer adjacent motif highlighted in bold ^23^.

### Protein expression and purification

Cas9 was purchased from IDT (Alt-R™ S.p. Cas9 Nuclease V3). The sequence of PEn with a C-terminal his tag was cloned into pET24a. The expression plasmid was then transformed into Escherichia coli BL21lDE3 Star (Thermofisher) for use in protein production. Autoinduction protocol^33^ was used for the over-production of PEn. Essentially, the culture was first grown overnight at 37 °C, before inoculation with 800 mL of ZYP autoinduction media, which was then grown at 37 °C with shaking until OD600 reached about 1-2. The temperature was then lowered to 18 °C and the culture was grown for a further 24 hours, before harvesting the cells by centrifugation. Cell pellets were stored at –80 °C until further use. The cell pellets were then resuspended in 20 mM HEPES, pH 7.5, 500 mM NaCl, 1 mM DTT, 10% glycerol, and lysed by one pass through an Emulsiflex C3 (Avestin). The lysate was clarified by centrifugation at 20,000 g for 20 minutes. The supernatant was supplemented with 10 mM imidazole and the lysate was loaded onto a 5 mL HiTrap column (Cytiva) equilibrated in the same buffer. The column was washed with 20 column volumes of 20 mM HEPES pH, 7.5, 500 mM NaCl, 1 mM DTT, 10% glycerol, 20 mM imidazole, before elution with 300 mM imidazole. The eluted protein was diluted to about 200 mM NaCl, before further purification on a 5 mL HiTrap Heparin SP column (Cytiva). Finally, the protein was further purified by size exclusion chromatography on a Superdex 200 (26/60) column (Cytiva) equilibrated in a buffer consisting of 20 mM HEPES, pH 7.5, 300 mM NaCl, 1 mM DTT, and 10% glycerol. The peak containing the PEn protein was pooled, concentrated to 10 mg/mL, flash-frozen in liquid nitrogen and stored at −80 °C until required.

### Plasmid transfection

K562 cells (10^6^ cells/condition) were transfected with 3.6 μg of PEmax or PEnmax and 1.2 μg of the pegRNA-containing plasmid using AMAXA Cell Line Nucleofector Kit V (VCA- 1003, Lonza) and U-16 program (Nucleofector 2b, Lonza). Cells transfected with TE buffer were used as negative controls.

### RNA transfection

K562 cells (2×10^5^ cells/condition) were electroporated with 2 μg of PEn mRNA and 100 pmol of synthetic pegRNA using the SF Cell Line 96-well Nucleofector™ Kit (Lonza) and the FF-120 pulse code (Nucleofector 4D; Lonza). Post-electroporation, cells were incubated at room temperature for 10 minutes without any disturbance and were thence transferred into pre-warmed medium in 96-well plates and incubated at 37°C with 5% CO_2_ for 72 hours before cell collection and DNA extraction. Cells transfected with TE buffer were used as negative controls.

### RNP and ssODN transfection

RNP complexes were assembled at room temperature using a final concentration of Cas9 at 3 μM, PEn at 2.7 μM and synthetic sgRNAs/pegRNAs at 5 μM. K562 cells (2×10^5^ cells/condition) were transfected with RNP complexes and ssODNs at 10 μM in the final transfection reaction, using the SF Cell Line 96-well Nucleofector™ Kit (Lonza), and the FF-120 pulse code (Nucleofector 4D) in the presence of a transfection enhancer (IDT). CD34^+^ HSPCs (2×10^5^ cells/condition) were transfected with RNP complexes and ssODNs at 10 μM in the final transfection reaction, using the P3 Primary Cell 4D-Nucleofector X Kit S (Lonza) and the CA-137 program (Nucleofector 4D) in the presence of a transfection enhancer (IDT). Post-electroporation, cells were incubated at room temperature for 10 minutes without any disturbance and were thence transferred into pre-warmed medium in 96-well plates and incubated at 37°C with 5% CO_2_ for 72 h for K562 cells or 6 days for HSPCs before cell collection and DNA extraction. Non-transfected cells or cells transfected with TE buffer or with enzyme only were used as negative controls.

### Small molecule compounds and drug treatment

DNA-PKc inhibitor AZD7648 was provided by AstraZeneca (Gothenburg, SE). PolΘ inhibitor, ART558 was purchased from MedChemExpress (HY-141520). All compounds were dissolved in dimethyl sulfoxide (DMSO) at a stock concentration of 10 mM. K562 cells were cultured for 24 h post-transfection in a medium containing 1 μM of AZD7648, or in combination with 3 μM ART558 (2i) when indicated. CD34^+^ cells were cultured in a medium containing 1 μM of AZD7648 and 3 μM ART558 (2i), for 24h post-transfection. All inhibitor treatments were followed by a cell wash and culture in control medium.

### HSPC differentiation

Transfected CD34^+^ HSPCs were differentiated into mature RBCs using a three-phase erythroid differentiation protocol, as previously described. During the first phase (day 0 to day 6), cells were cultured in a basal erythroid medium supplemented with 100 ng/ml recombinant human SCF (PeproTech), 5 ng/ml recombinant human IL3 (PeproTech), 3 IU/ml EPO Eprex (Janssen-Cilag) and 10^−6^M hydrocortisone (Sigma). During the second phase (day 6 to day 9), cells were co-cultured with MS5 stromal cells in the basal erythroid medium supplemented with 3 IU/ml EPO Eprex (Janssen-Cilag). During the third phase (day 9 to day 20), cells were co-cultured with stromal MS5 cells in a basal erythroid medium without cytokines. Heat-inactivated human AB serum was added during the third phase of the differentiation (10%; day 13 to day 20). Erythroid differentiation was monitored by flow cytometry analysis of CD36, CD71, GPA, Band3, and α4-Integrin erythroid surface markers and of enucleated cells using the DRAQ5 double-stranded DNA dye. 7AAD was used to identify live cells.

### Colony-forming cell (CFC) assay

CD34^+^ HSPCs were plated at a concentration of 500 cells/mL in a methylcellulose-based medium (GFH4435, Stem Cell Technologies) under conditions supporting erythroid and granulocyte/monocyte differentiation. BFU-E and CFU-GM colonies were counted after 14 days. Colonies were randomly picked and collected as bulk populations (*n*=25 colonies) to evaluate the editing efficiency, globin expression by RT-qPCR and RP-HPLC. BFU-Es were randomly picked and collected as single colonies (around 24 colonies per sample) to evaluate the efficiency and globin expression by RT-qPCR.

### Genomic DNA extraction and sequencing for on- and off-target analysis

Genomic DNA from K562 cells was extracted using Quick Extract solution (LGC Biosearch Technologies) or the PureLink Genomic DNA Mini Kit (Invitrogen), 3 days post-transfection, following manufacturer’s instructions. Genomic DNA from CD34^+^ was harvested 6 days post-transfection using the PureLink Genomic DNA Mini Kit (Invitrogen) following the manufacturer’s instructions.

To evaluate the editing efficiency in K562 cells’ screening experiments (**Figure S1A**), *HBG1/2* promoters’ on-target sites were PCR-amplified, with 5’-AAAAACGGCTGACAAAAGAAGTCCTGGTAT-3’ forward primer and 5’-ATAACCTCAGACGTTCCAGAAGCGAGTGTG-3’ reverse primer, using the Recombinant Taq DNA Polymerase (Thermo Fisher) according to the manufacturer’s instructions, and subjected to Sanger sequencing. The precise prime edits, i.e. expected insertion size, and the Indels were evaluated using the TIDE software (http://shinyapps.datacurators.nl/tide/).

The *HBG* promoters were amplified with 5’-GGAATGACTGAATCGGAACAAGG-3’ forward primer and 5’-CTGGCCTCACTGGATACTCT-3’ reverse primer for pegRNA-115 and sgRNA-115 targeting samples. The hybrid *HBG1/2* promoter was amplified by a nested PCR with 5’-GTTTTAAAACAACAAAAATGAGGGAAAGA-3’ forward primer and 5’-GTTGCTTTATAGGATTTTTCACTACAC-3’ reverse primer using Phusion Flash High-Fidelity 2x Mastermix (F548, Thermo Scientific) in a 30 μL reaction, containing 1.0 μL of genomic DNA extract and 0.2 μM of target-specific primers (98 °C for 3 min, followed by 40 cycles of 98 °C for 10 seconds, 60 °C for 5 seconds, and 72 °C for 18 seconds), followed by amplification using pegRNA-115 and sgRNA-115 targeting sample primers. Amplicons were generated with Phusion Flash High-Fidelity 2x Mastermix (F548, Thermo Scientific) in a 15 μL reaction, containing 1.5 μL of genomic DNA extract or 1^st^ PCR product (for nested PCR) and 0.2 μM of target-specific primers with barcodes and NGS adapters. PCR cycling conditions for Phusion Flash High-Fidelity 2x Mastermix were: 98 °C for 3 min, followed by 30 cycles of 98 °C for 10 seconds, 60 °C for 20 seconds, and 72 °C for 30 seconds and final elongation at 72 °C for 2 min. All amplicons were purified using HighPrep PCR Clean-up System (MagBio Genomics). Size, purity, and concentration of amplicons were determined using a fragment analyzer (Agilent). To add Illumina indexes to the amplicons, samples were subjected to a second round of PCR. Indexing PCR was performed using KAPA HiFi HotStart Ready Mix (Roche), 0.067 ng of PCR template and 0.5 µM of indexed primers in the total reaction volume of 25 µL. PCR cycling conditions were 72 °C for 3 minutes, 98 °C for 30 seconds, followed by 10 cycles of 98 °C for 10 seconds, 63 °C for 30 seconds, and 72 °C for 3 min, with a final extension at 72 °C for 5 minutes. Samples were purified with the HighPrep PCR Clean-up System (MagBio Genomics) and analyzed using a fragment analyzer (Agilent). Samples were quantified using a Qubit 4 Fluorometer (Life Technologies) and subjected to sequencing using Illumina NextSeq system according to manufacturer’s instructions. **Table S3** lists the primers used for on- and off-target analysis by deep sequencing.

### Bioinformatic analysis

Demultiplexing of the NGS sequencing data was performed using bcl2fastq software. The fastq files were analyzed using CRISPResso2 V2.2.12 in the HDR or prime editing mode. Detailed parameters are listed in **Table S4**.

### RT-qPCR

Total RNA was extracted from BFU-E pools and erythroblasts (day 13) using the RNeasy micro kit (QIAGEN), and single colonies using the AllPrep DNA/RNA kit (QIAGEN). RNA was treated with DNase using the DNase I kit (Invitrogen), following the manufacturer’s instructions. Mature transcripts were reverse-transcribed using SuperScript III First Strand Synthesis System for RT-qPCR (Invitrogen) with oligo (dT) primers. RT-qPCR was performed using the iTaq universal SYBR Green master mix (Bio-rad) and the CFX384 Touch Real-Time PCR Detection System (Biorad). **Table S5** lists the primers used for RT-qPCR analysis.

### Evaluation of the 4.9-kb deletion

Digital Droplet PCR (ddPCR) was performed using a primer/probe mix (BioRad) to quantify the frequency of the 4.9-kb deletion. Control primers annealing to *HBG2* were used as DNA loading control. ddPCR was performed using the ddPCR supermix (no dUTP) to quantify the frequency of *HBG2* loss over *HBG1* in each treated sample. Data were acquired through QX200 analyzer (Biorad) and results were analyzed with QuantasoftTM Analysis Pro (Biorad). **Table S6** lists the primers used for the ddPCR analysis.

### Flow cytometry analysis

HSPC-derived erythroid cells were stained with an antibody recognizing GPA erythroid surface marker (1/100 PE-Cy7 conjugated anti-GPA antibody, 563666, BD Biosciences), then fixed with 0.05% cold glutaraldehyde and permeabilized with 0.1% TRITON X-100. After fixation and permeabilization, cells were stained with either an antibody recognizing HbF (1/5 FITC-conjugated anti-HbF antibody, clone 2D12 552829 BD Biosciences), or an antibody recognizing HbS (1/20 anti-HbS antibody, H04181601, BioMedomics) followed by the staining with a secondary antibody recognizing rabbit IgG (1/200 BV421-conjugated anti-rabbit IgG, 565014, BD Biosciences). Flow cytometry analysis of CD36, CD71, GPA, BAND3 and α4-Integrin erythroid surface markers was performed using a V450-conjugated anti-CD36 antibody (1/100 561535, BD Horizon), a FITC-conjugated anti-CD71 antibody (1/100 555536, BD Biosciences), a PE-Cy7 conjugated anti-GPA antibody (1/100 563666, BD Biosciences), a PE-conjugated anti-BAND3 antibody (1/50 9439, IBGRL) and an APC-conjugated anti-CD49d antibody (1/20 559881, BD Biosciences). Flow cytometry analysis of enucleated or viable cells was performed using double-stranded DNA dyes (DRAQ5, 65-0880-96, Invitrogen and 7AAD, 559925, BD, respectively). Flow cytometry analyses were performed using Novocyte flow cytometers (Agilent Technologies). Data were analyzed using the FlowJo (BD Biosciences) software.

### Statistics

Data visualization and statistical analysis were conducted using GraphPad Prism 9 (GraphPad Software, Inc.). Figure legends contain information on statistical tests, sample sizes, and P values. Briefly, we used Shapiro-Wilk test to evaluate if data follow a Gaussian distribution. For comparison between two groups, we used the parametric unpaired t-test if data followed a Gaussian distribution and the non-parametric Mann-Whitney test if data did not follow a Gaussian distribution. No data were excluded from the analyses. No statistical method was used to predetermine sample size. The experiments were not randomized. The investigators were not blinded during experiments and outcome assessment

## Data availability statement

All data supporting the findings of this study are available within the paper and its Supplementary Information.

## Acknowledgments

We thank Dr. Sandra Manceau for the collection of the blood samples and the technological core facilities of the SFR Necker (Inserm US24 / CNRS UAR3633) including the flow cytometry platform and the imaging facility. This work was supported by state funding from the French National Research Agency (Agence Nationale de la Recherche; ANR-10-IAHU-01 and ANR-22-CE17-0028 PEMGeT), the European Commission (HORIZON-PathFinder EdiGenT grant no. 101070903), the COST (European Cooperation in Science and Technology (the COST Action Gene Editing for the treatment of Human Diseases, CA21113) and ERDERA, which has received funding from the European Union’s Horizon Europe research and innovation programme under grant agreement N°101156595. This study is part of the Université Paris Cité IdEx #ANR-18-IDEX-0001 funded by the French Government through its “Investments for the Future” program.

## Author contributions

A.C. Conceptualization, Formal analysis, Investigation, Writing – original draft; P.A. Conceptualization, Formal analysis, Investigation, Writing – original draft; M.B.D. Formal analysis, Investigation; M.F. Data curation Formal analysis, Investigation; M.P. Formal analysis, Investigation; M.M. Conceptualization, Funding acquisition, Supervision, Validation – Verification, Writing – review & editing; M.B. Conceptualization, Project administration, Supervision, Validation – Verification, Writing – original draft Writing – review & editing; A.M. Conceptualization, Funding acquisition, Project administration, Supervision, Validation – Verification, Writing – review & editing.

