## supplemental information for "Enhanced γ-globin reactivation and sickle cell correction through a repressor-to-activator motif switch in the *HBG1/2* promoters"

**SUPPLEMENTARY TABLES**

Table S1: List of pegRNAs and springRNAs sequences used in the study
Table S2: List of ssODNs sequences targeting the -115 region of the HBG promoters
Table S3: List of primers used for on- and off-target analysis by deep sequencing
Table S4: Crispresso2 analysis
Table S5: List of primers used for RT-qPCR analysis
Table S6: Primers and probe used for the 4.9-kb deletion ddPCR analysis

### SUPPLEMENTARY FIGURE LEGENDS

#### Figure S1

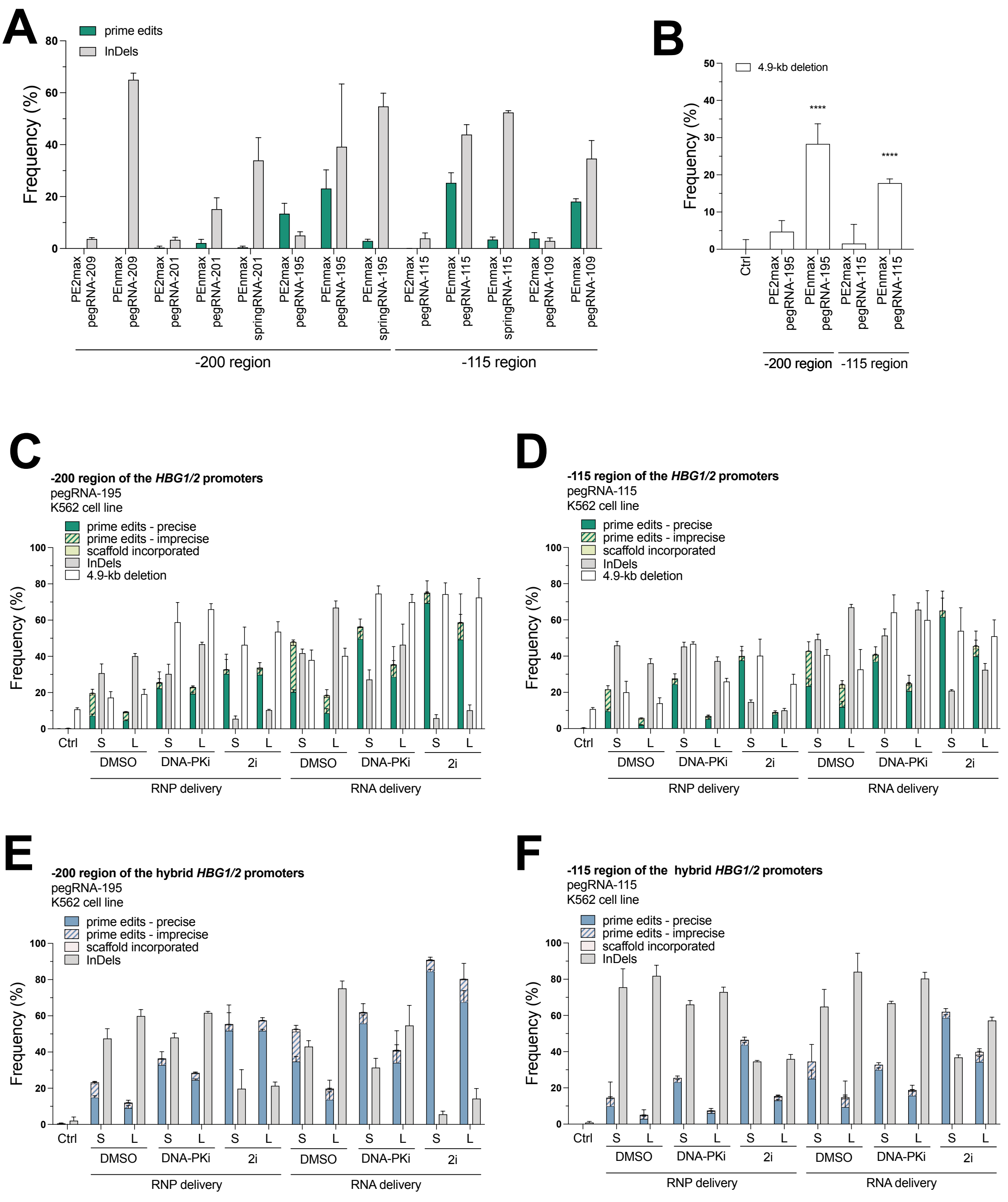


**Figure S1: Design and screening of pegRNAs and springRNAs targeting the *HBG1/2* promoters in K562 cells.**

(A) Frequency of the desired insertion (prime edits) and InDels in the *HBG1/2* promoters induced by pegRNAs or the springRNAs in combination with PEnmax or PE2max in K562 cells. The *HBG1/2* promoters were amplified and subjected to Sanger sequencing. Data were analyzed using the TIDE software. Bars represent the mean ± SD of 3 biologically independent replicates.

(B) The frequency of the 4.9-kb deletion was measured by ddPCR in control (Ctrl, cells transfected with Tris-EDTA, TE) and edited cells.

(C, D) Percentage of total *HBG1/2* promoters containing the total prime edits (precise, imprecise, scaffold incorporated) and the InDels induced by (C) pegRNA-195_S or pegRNA-115_L, or (D) pegRNA-115_S or pegRNA-115_L, delivered as RNA or RNP in K562 cells treated or not with DNA-PKi or DNA-PKi and Polθi inhibitors. The *HBG1/2* promoters were amplified and subjected to NGS. Data were analyzed with the CRISPResso2 webtool. We report also the frequency of the 4.9-kb deletion measured by ddPCR for control (ctrl, TE-transfected cells) and prime-edited conditions. Bars represent the mean ± SD of 3 technical replicates.
(E, F) Percentage of the hybrid *HBG1/2* promoters containing the prime edits (precise, imprecise, scaffold incorporated) and InDels induced by (E) pegRNA-195_S or pegRNA-115_L or (F) pegRNA-115_S or pegRNA-115_L, delivered as RNA or RNP in K562 cells treated or not with DNA-PKi or DNA-PKi and Polθi inhibitors. The hybrid *HBG1/2* promoters were amplified and subjected to NGS. Data were analyzed with the CRISPResso2 webtool. Bars represent the mean ± SD of 3 biological replicates.

**Figure S2**

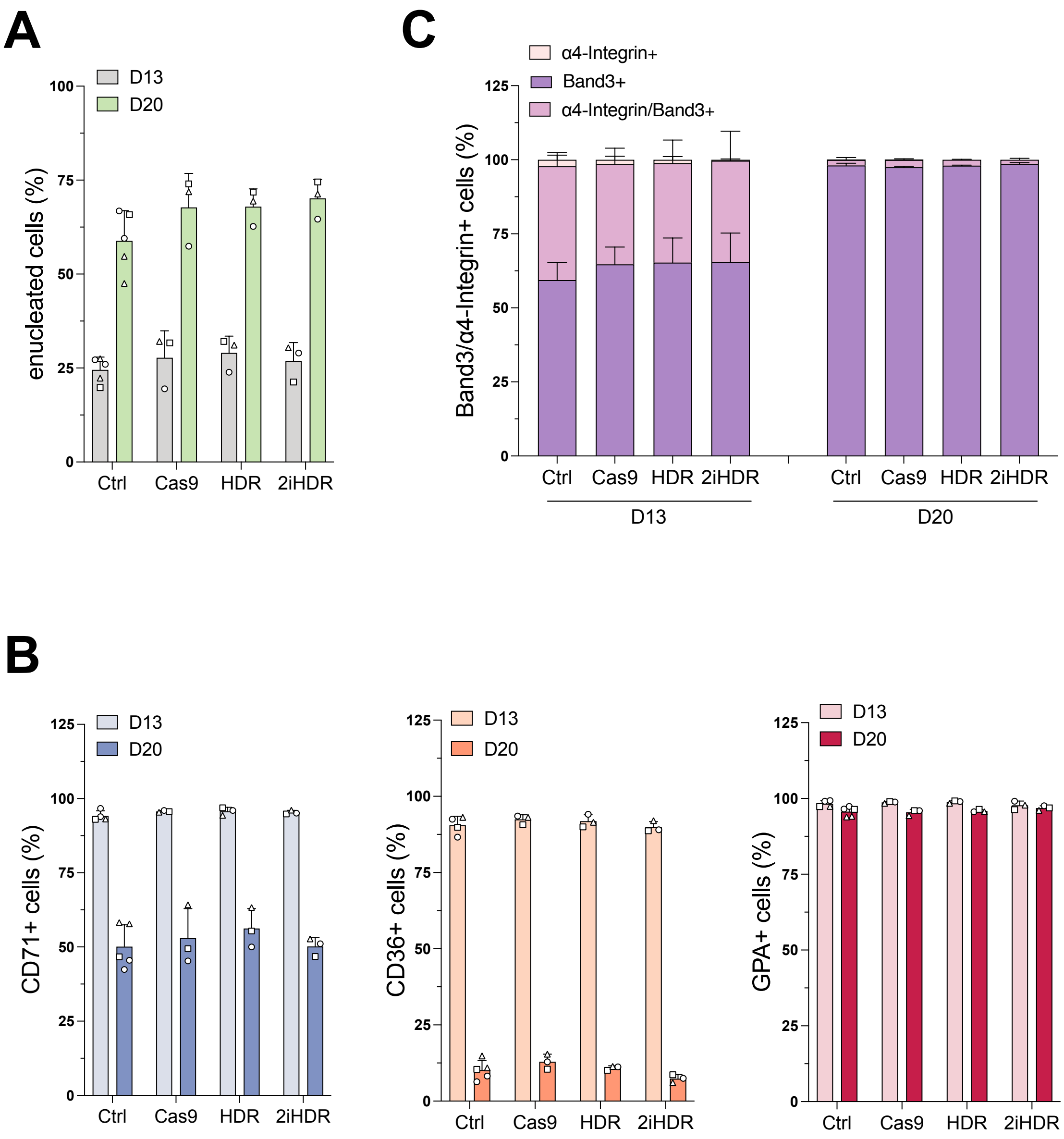


**Figure S2: The Cas9-based strategies do not impact the erythroid differentiation**
(A) Frequency of enucleated cells at day 13 and 20 of erythroid differentiation, as measured by flow cytometry analysis after DRAQ5 nuclear staining in control and edited samples. Data are expressed as mean ± SD (n=3 biologically independent experiments; 2 different donors). No statistical differences were observed between control and edited samples using two-way ANOVA with Tukey's multiple comparisons test.
(B) Frequency of (left) CD71+, (middle) CD36+ and (right) GPA+ cells at day 13 and 20 of the erythroid differentiation, as measured by flow cytometry analysis of CD36, CD71, and GPA erythroid markers in control and edited samples. Data are expressed as mean ± SD (n=3 biologically independent experiments; 2 different donors). No statistical differences were observed between control and edited samples using two-way ANOVA with Tukey's multiple comparisons test.

(C) Frequency of α4-Integrin+, Band3+ and α4-Integrin+/Band3+ in 7AAD-/GPA+ cells at day 13 and 20 of erythroid differentiation, as measured by flow cytometry analysis of α4-Integrin and Band3 erythroid markers in control and edited samples. Data are expressed as mean ± SD (n=3 biologically independent experiments; 2 different donors). No statistical differences were observed between control and edited samples using two-way ANOVA with Tukey's multiple comparisons test.

### Figures S3

A

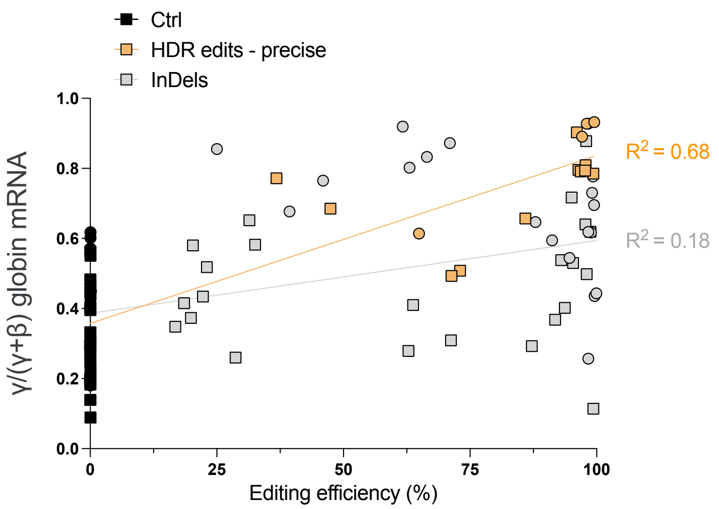


**Figure S3: Cas9-HDR-edited BFU-E express high HbF levels.**
(A) Correlation between γ-globin mRNA expression and editing efficiency in single erythroid BFU-E (n=32 to 74 single BFU-E for each group). γ-globin mRNA expression was normalized to α-globin mRNA and expressed as a percentage of the total β- and γ- globin mRNA. For BFU-E containing the HDR precise edits, $R^2$ = 0.68, Y = 0,004792*X + 0,3577; for BFU-E containing InDels, $R^2$ = 0.18, Y = 0,002081*X + 0,3862 (simple linear regression).
